# A novel reporter gene assay for pyrogen detection

**DOI:** 10.1101/613190

**Authors:** Qing He, Chuan-fei Yu, Lang Wang, Yong-bo Ni, Heng Zhang, Ying Du, Hua Gao, Jun-zhi Wang

**Affiliations:** National Institutes for Food and Drug Control, Beijing, China

**Author notes:** Corresponding author: Jun-zhi Wang, Hua Gao, National Institutes for Food and Drug Control, No. 31, Huatuo Road, Daxing District, Beijing 102629, China. contributed equally (Qing He, Chuan-fei Yu, Lang Wang).

**Keywords:** Fever, Pyrogens, Nuclear factor kappa B, RAW 264.7, Lipopolysaccharide, Lipoteichoic acid, Zymosan

## Abstract

Fever is a systemic inflammatory response of the body to pyrogens. Nuclear factor κB (NF-κB) is a central signalling molecule that causes the excessive secretion of various proinflammatory factors induced by pyrogens. This study explored the feasibility of a novel reporter gene assay (RGA) for pyrogen detection using RAW 264.7 cells stably transfected with the NF-κB reporter gene as a pyrogenic marker. Pyrogen was incubated with the transgenic cells, and the intensity of the fluorescence signal generated by luciferase secreted by the reporter gene was used to reflect the degree of activation of NF-κB, so as to quantitatively detect the pyrogens. The RGA could detect different types of pyrogens, including the lipopolysaccharide (LPS) of gram-negative bacteria, the lipoteichoic acid (LTA) of gram-positive bacteria, and the zymosan of fungi, and a good dose-effect relationship was observed in terms of NF-κB activity. The limits of detection of the RGA to those pyrogens were 0.03 EU/ml, 0.001 μg/ml, and 1 μg/ml, respectively. The method had good precision and accuracy and could be applied to many biological products (e.g., nivolumab, rituximab, bevacizumab, etanercept, basiliximab, *haemophilus influenzae* type b conjugate vaccine, 23-valent pneumococcal polysaccharide vaccine, and group A and group C meningococcal conjugate vaccine). The results of this study suggest that the novel RGA has a wide pyrogen detection spectrum and is sufficiently sensitive, stable, and accurate for various applications.

**Importance:** Pyrogen testing is mandatory and a critical method to ensure the safety of parenteral products including vaccines.

Currently, only two pharmacological tests, including the rabbit pyrogen test and the bacterial endotoxins test (BET), are applied to evaluate pyrogenic contamination in parenteral pharmaceuticals by most of state pharmacopoeias. Although generally reliable, both of these assays have shortcomings. The rabbit test is not quantitative but is expensive and involves the use of animals. It can also produce varying responses depending on the strain, age and housing conditions of the rabbits. The BET, however, does not detect pyrogens other than gram-negative bacterial endotoxins and is often problematic when used to test solutions with a high protein content.

To overcome these shortcomings and satisfy the growing need for new methods prompted by the constantly increasing production of biological compounds, it is necessary to develop the novel assay for pyrogen detection.

**Highlights:**

- This novel reporter gene assay can detect different types of pyrogens, including the lipopolysaccharide of gram-negative bacteria, the lipoteichoic acid of gram-positive bacteria, and the zymosan of fungi.
- The novel reporter gene assay is sufficiently sensitive, stable, and accurate for various applications.

## 1. Introduction

Pyrogens are fever-inducing substances, including exogenous pyrogens [e.g., the lipopolysaccharide (LPS) of gram-negative bacteria, the lipoteichoic acid (LTA) of gram-positive bacteria, the peptidoglycan (PGN) and lipoprotein (LP) of gram-negative/positive bacteria, and the zymosan of fungi] and endogenous pyrogens (e.g., steroids, prostaglandin E, and proinflammatory cytokines) [1,2]. Pyrogen testing is mandatory and a critical method to ensure the safety of parenteral products. The Chinese Pharmacopoeia (CP) has adopted the rabbit pyrogen test (RPT) and the bacterial endotoxin test (BET) for detecting pyrogenic contamination in products [3]. Our laboratory consumes approximately 1000-1500 rabbits per year for the RPT, along with large quantities of manpower and material resources. Nevertheless, the results of the RPT are often affected by many factors, such as environmental, animal, and operating factors. Our laboratory also consumes approximately 15,000-20,000 vials of horseshoe crab reagents per year for the BET, which can only detect the LPS of gram-negative bacteria. Most domestic horseshoe crabs are usually not reused to produce reagents; meanwhile, horseshoe crab populations are becoming increasingly scarce due to unreasonable capture practices, habitat loss, and pollution. The European Pharmacopoeia (EP) has adopted the monocyte activation test (MAT), which is mainly based on the use of monocytes and macrophages involved in fever and proinflammatory cytokines [e.g., interleukin (IL)-6, IL-1β, and tumor necrosis factor (TNF)-α] as pyrogenic markers, to replace those traditional pyrogen tests [4]. The MAT often needs large amounts of human blood and its convenience needs to be improved. The representativeness of using a single proinflammatory cytokine as the pyrogenic marker is also limited in theory; however it does not involve the use of animals *in vivo*, has a wide pyrogen detection spectrum, and follows the 3Rs principle [5,6], which has gained widespread attention from researchers.

In essence, fever is a systemic inflammatory response of the body to pyrogens [7-9]. However, pyrogens can stimulate the body through different mechanisms and can induce the excessive production of different proinflammatory factors, such as ILs, TNF-α, CC chemokine ligand 5, CXC chemokine ligand 1, and prostaglandins, from monocytes and macrophages [10-13]. For example, LPS mainly binds CD14 and Toll-like receptor 4 (TLR4), and activated TLR4 can promote inflammation mainly via pathways dependent on MyD88 (MyD88→NF-κB and IRF-5→synthesis of proinflammatory cytokines, including TNF-α, IL-1β, and IL-6) and TIR-domain-containing adapter-inducing interferon β (TRIF) (TRIF→NF-κB) [14-18]. LTA mainly binds TLR2 to activate NF-κB, resulting in the production of proinflammatory mediators, such as TNF-α, IL-1, IL-6, IL-8, nitric oxide (NO), and chemokines [19-22]. LP, diacylated LP, and triacylated LP mainly bind TLR2, a dimer formed with TLR2 and TLR6, and a dimer formed with TLR2 and TLR1. Activated TLRs can promote inflammation mainly caused via the MyD88-dependent pathway (MyD88→NF-κB/MAPK→synthesis of proinflammatory cytokines) [23,24]. Zymosan can bind TLR2 to activate NF-κB, resulting in the production of proinflammatory mediators, such as TNF-α, IL-1β, and IL-8 [25-28]. The mechanisms of pyrogens stimulating the secretion of proinflammatory factors in the body often involve the activation of NF-κB, which is the central signalling molecule mediating the inflammatory response [29-32]. Thus, it is reasonable to use NF-κB as a representative pyrogenic marker. Therefore, the main aim of the present study was to evaluate the feasibility of utilizing murine macrophage RAW 264.7 cells transfected with the NF-κB reporter gene to detect pyrogens.

## 2. Experiments

### 2.1. Materials and methods

#### 2.1.1. Reagents

The national standard for bacterial endotoxins is LPS, which was obtained from *Escherichia coli O55:B5* [10000 endotoxin units (EU)/vial, batch 150600-200707, identical to the 2nd international WHO standard for endotoxin 94/580 from *Escherichia coli* O113:H10] and was provided by the National Institutes for Food and Drug Control (NIFDC). The following materials were also used in this work: LTA (Sigma-Aldrich, Cat # L3265), zymosan (Sigma-Aldrich, Cat # Z4250), foetal bovine serum (FBS, Gemini, Cat # 900-108), penicillin-streptomycin (Gibco, Cat # 15140-122), L-glutamine (Gibco, Cat # 25030-081), hygromycin B (Amresco, Cat # V900372), Bright-Glo Luciferase Assay reagent (Promega, Cat # E2650), phosphate-buffered saline (PBS, HyClone, Cat # SH30256.01), trypsin-EDTA (Gibco, Cat # 25200-056), DMEM (Gibco, Cat # 11995-065), pyrogen-free water for the BET (Zhanjiang A&C Biological, LTD), *Tachypleus* amebocyte lysate (TAL, Zhanjiang A&C Biological, LTD), nivolumab injection reagent (Bristol-Myers Squibb Holdings Pharma, LTD Liability Company), rituximab injection reagent (Roche Diagnostics, GmbH), bevacizumab injection reagent (Roche Diagnostics, GmbH), etanercept solution for injection (Pfizer Ireland Pharmaceuticals), *haemophilus influenzae* type b conjugate vaccine (Yuxi Walvax Biotechnology Co., LTD), 23-valent pneumococcal polysaccharide vaccine (Yuxi Walvax Biotechnology Co., LTD), group A and group C meningococcal conjugate vaccine (Yuxi Walvax Biotechnology Co., LTD), and basiliximab for injection (Novartis Pharma Stein AG).

#### 2.1.2. Consumables

Ninety-six-well plates were used for both the RGA (Corning, Cat # 3917, white, flat bottom, tissue culture treated, polystyrene) and the BET (Corning, Cat # M9005, flat bottom, polystyrene, tissue culture-treated). Mouse IL-1β, IL-6 and TNF-α ELISAs were performed using commercially available kits (Xin Bo Sheng Co.). Other reagents/materials were purchased as sterile and free of pyrogens, and glassware was baked at 250°C for 1 h.

#### 2.1.3. Construction of the reporter gene vector

The pCM1.1_luc_hygro vector contains a minimal promoter followed by a luciferase gene. NF-κB response element (5_-TCCTCGGAAAGTCCCCTCTGAGATCCTCGGAAAGTCCCCTCTGAGATCTCAGAGGGGACTTTCCGAGGA-3_) was synthesized by overlap PCR and inserted into the multiple cloning site ahead of the mini-promoter region, and the positive clone was verified by DNA sequencing.

#### 2.1.4. Development of RAW 264.7 cells stably transfected with the reporter gene vector

The plasmid pCM1.1_ NF-κB _luc_hygro was introduced into RAW 264.7 cells (ATCC) by electroporation. The cells were selected at 48 h after transfection in selective media (DMEM containing 10% FBS, 1% penicillin-streptomycin, 1% glutamine, and 150 μg/ml hygromycin B). After being selected for 3 weeks, hygromycin-resistant cells were then cloned by limited dilution to obtain a single cell clone and were then screened for the induction of luciferase activity by treatment with gradient concentrations of LPS (e.g. 1000 ng/ml, 100 ng/ml, and then 1:3 dilutions, 10 series). The resulting positive clones were routinely maintained in the selective media.

#### 2.1.5. FACS analysis

RAW 264.7 cells were centrifuged (300 g×5 min) at 4°C, washed twice with ice-cold PBS, and then blocked with 200 μg/ml mouse IgG (Jackson ImmunoResearch, Cat # 015-000-003) on ice for 10-20 min. The cells were resuspended in PBS containing 4% bovine calf serum (BCS) at a concentration of 2**×**10^6^ cells/ml and were aliquoted into 96-well plates (50 μl/well). Then, 50 μl of 2 μg/ml fluorescence-labelled antibodies was added for the detection of TLR2 (R&D, Cat # FAB1530G), TLR4 (R&D, Cat # FAB27591G), and TLR6 (R&D, Cat # FAB1533G); rat IgG2a Alexa Fluor (AF) 488-conjugated antibody was used as an isotype control (R&D, Cat # IC006G). The cell-antibody mixture was incubated on ice for 45 min in the dark, washed twice with PBS containing 4% BCS, and resuspended in 200 μl of 5 μg/ml propidium iodide (PI, Sigma-Aldrich, Cat # P4170) in PBS to stain dead cells. Data were collected on a BD FACSCanto system and analysed using FlowJo software.

#### 2.1.6. RGA

Selected cells in the logarithmic growth stage were washed with PBS and digested with 0.25% trypsin-EDTA. Confluent cell monolayers were suspended in the selective medium at the required concentration, seeded into 96-well plates (100 μl/well), and allowed to attach for 24 h. Then, the selective medium was discarded, and sample solutions prepared with assay medium (DMEM containing 10% FBS) were added to the 96-well plates (100 μl/well, *n*=4). The plates were incubated at 37°C in an atmosphere of 5% CO_2_ in air for a period. After incubation, Bright-Glo Luciferase Assay reagent was added to the 96-well plates (100 μl/well), which were subsequently shaken for 1 min. Finally, luciferase activity was determined using a Luminoskan Ascent reader. If necessary, commercial ELISA kits were used to detect the levels of proinflammatory factors (e.g., IL-1β, IL-6, and TNF-α) in the supernatants.

#### 2.1.7. BET

The kinetic chromogenic TAL assay was performed according to the manufacturer’s instructions (Zhanjiang A&C Biological, LTD). One hundred microliters/well of the sample/standard solutions (*n*=2) prepared with pyrogen-free water was mixed with an equal volume of an endotoxin-specific TAL reagent, also prepared with pyrogen-free water, in 96-well plates. The rate of colour development was measured at 37°C using a specially equipped microplate reader (Synergy HT, BioTek Instruments, Inc.). The endotoxin contents of the samples were calculated according to the parallel line assay method using the logarithmically transformed dose and the rate of colour development and are expressed as EU/ml, referring to the standard endotoxin.

### 2.2. Statistical analysis

All experiments were repeated three times. The data are expressed as the mean and standard error of the mean (SEM). The data were compared between groups using the Student’s *t*-test.

## 3. Results

### 3.1. Identification of TLR2, TLR4, and TLR6 on RAW 264.7 cells

The results of this experiment are presented in Figure 1. The data show that the RAW 264.7 cells expressed the main receptors that can bind to pyrogens, including TLR2, TLR4, and TLR6.

**Fig. 1.**
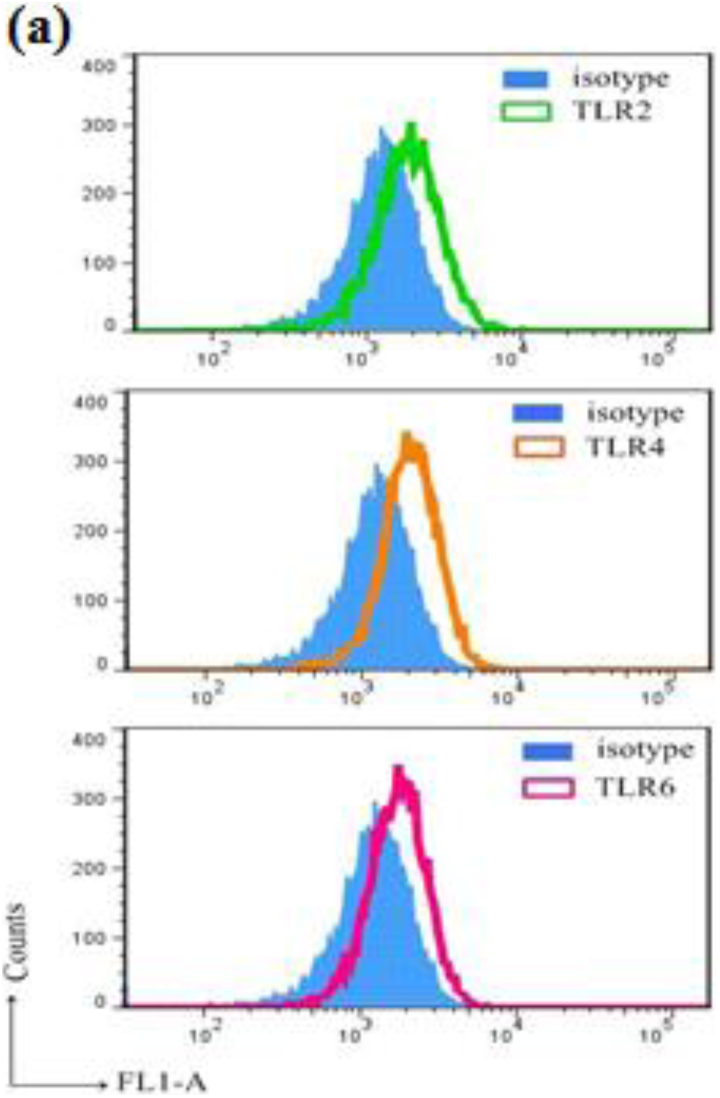
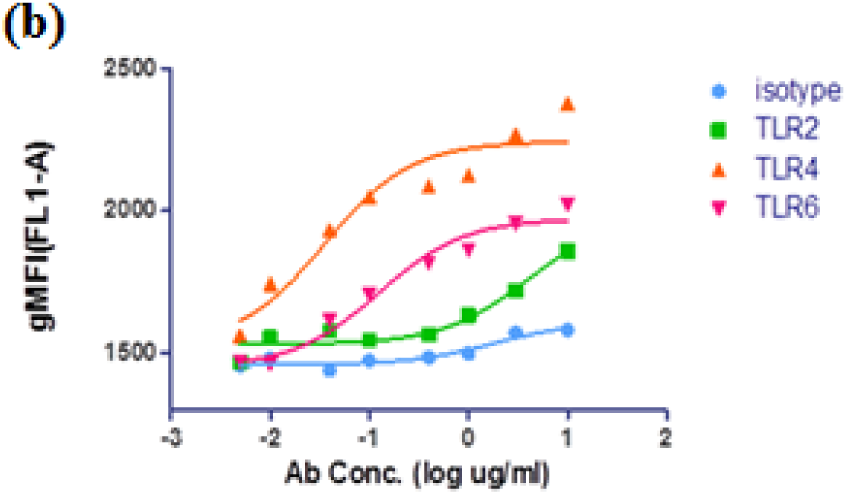
Expression of TLR2 (green line), TLR4 (orange line), and TLR6 (purple line) on RAW 264.7 cells (a) and binding curves of the corresponding antibodies (Ab, concentrations from 10 μg/ml to 0.005 μg/ml, at 1:3 dilutions) to TLR2, TLR4, and TLR6 on RAW 264.7 cells (b) were analysed by FACS. Rat IgG2a AF488 (blue shadow or line) was used as an isotype control.

### 3.2. Correlation between NF-κB activation and proinflammatory factor secretion

The results of this experiment are presented in Figure 2. The Pearson’s correlation coefficients between the activation of NF-κB and the secretion of proinflammatory factors (e.g., IL-1β, IL-6, and TNF-α) were 0.967, 0.895, and 0.721, respectively, for LPS; 0.836, 0.986, and 0.915, respectively, for LTA; and 0.981, 0.950, and 0.838, respectively, for zymosan. The data show that the dose-effect trends of the LPS-, LTA-, and zymosan-induced NF-κB activation were consistent with those of the secretion of proinflammatory factors induced by those pyrogens, suggesting that a good correlation between the activation of NF-κB and the secretion of proinflammatory factors.

**Fig. 2.**
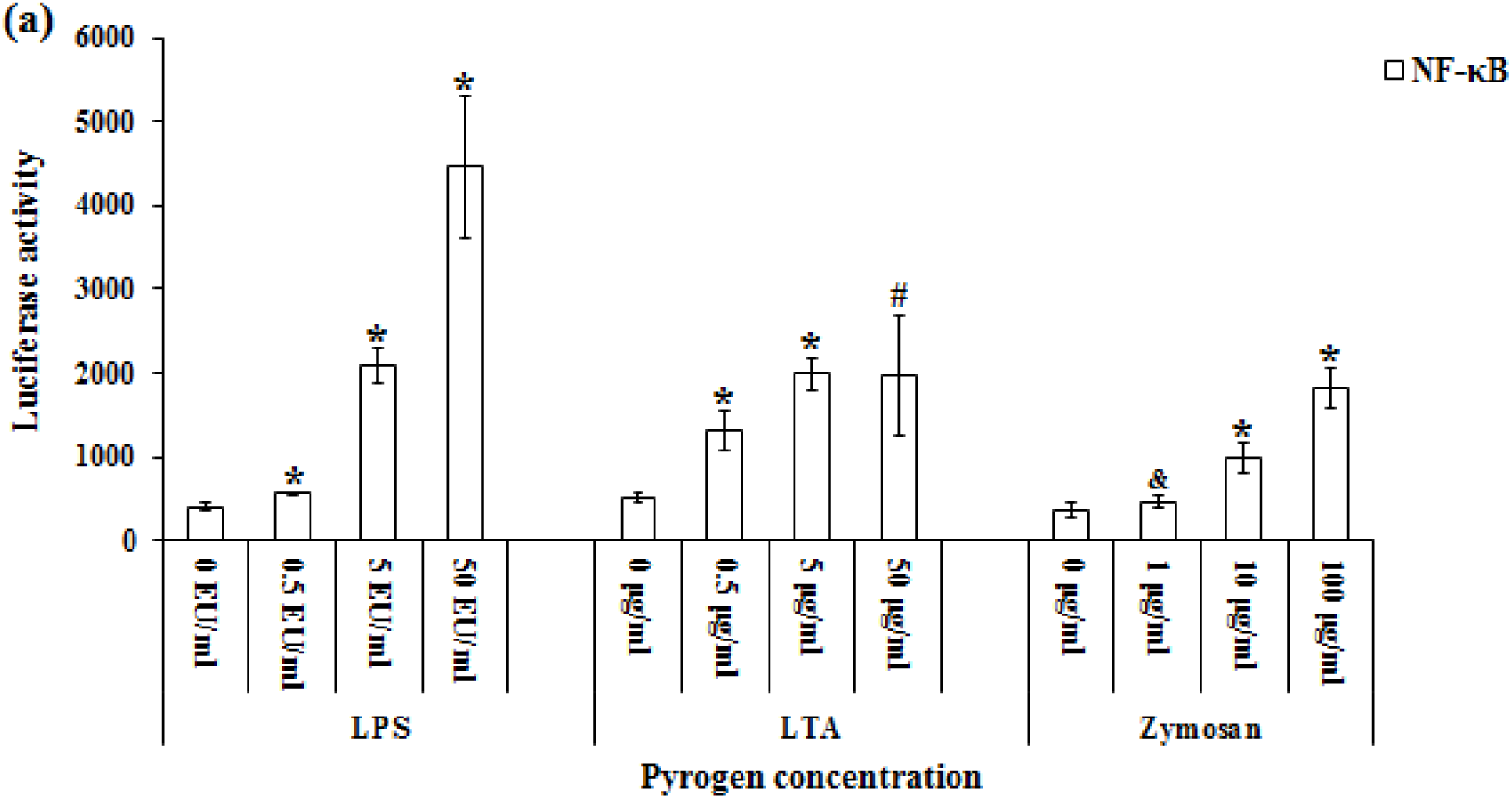
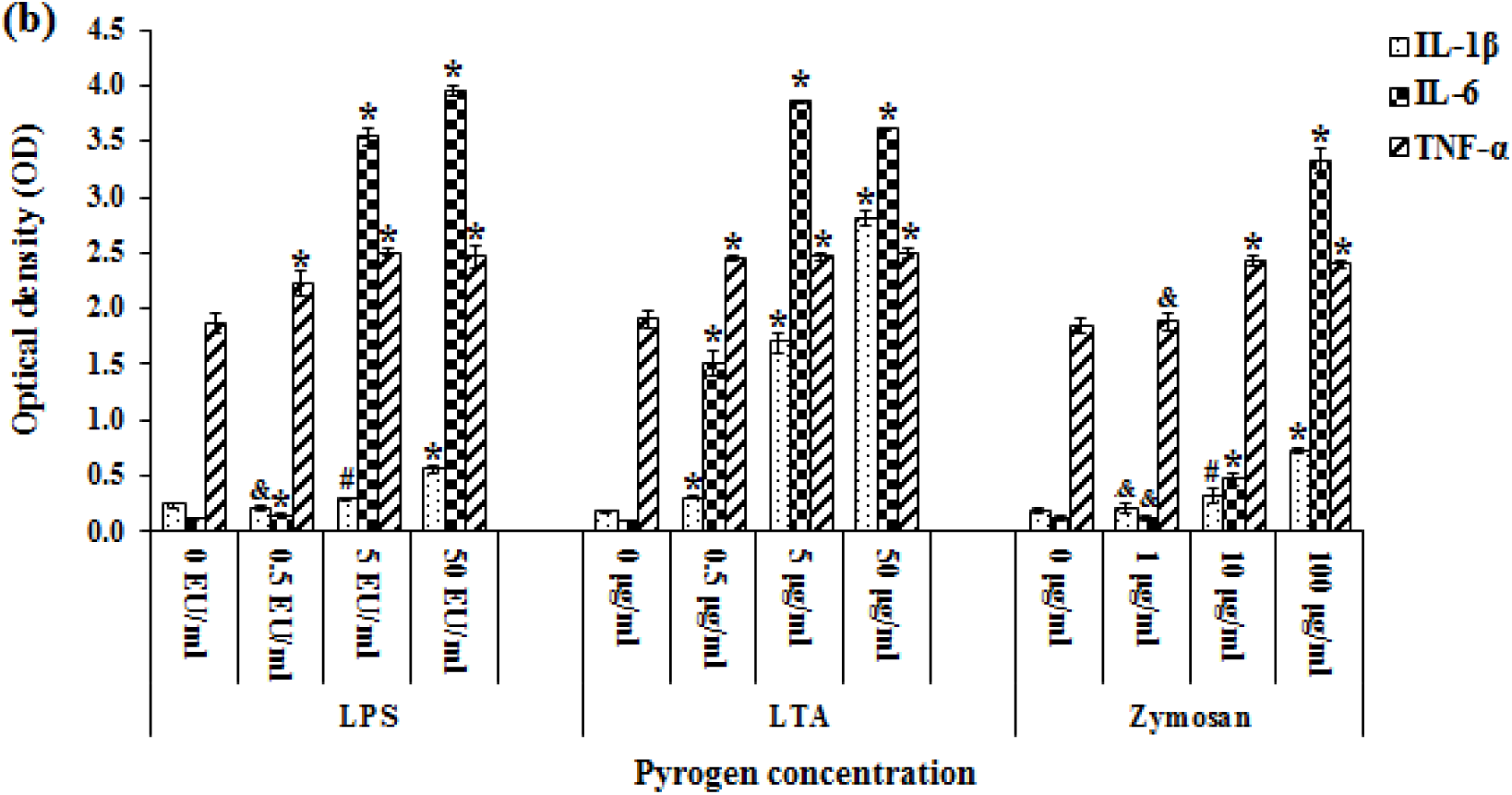
The activity of NF-κB (a) and the secretion of the proinflammatory factors (b) IL-1β, IL-6, and TNF-α in RAW 264.7 cells at a density of 10×10^5^ cells/ml after stimulation with LPS, LTA, and zymosan for 24 h (*n*=4). & *P*>0.05 vs. the negative control (e.g., 0 EU/ml, 0 μg/ml) # *P*<0.05 vs. the negative control * *P*<0.01 vs. the negative control

### 3.3. The time-effect relationships of pyrogens (LPS, LTA, and zymosan) activating NF-κB

The results of this experiment are presented in Figure 3. The data show that the different pyrogens, i.e., LPS, LTA, and zymosan, could activate NF-κB in a dose-dependent manner. Increasing the stimulation time up to 10-12 h, almost all concentrations of the three pyrogens could activate NF-κB to an extreme extent.

**Fig. 3.**
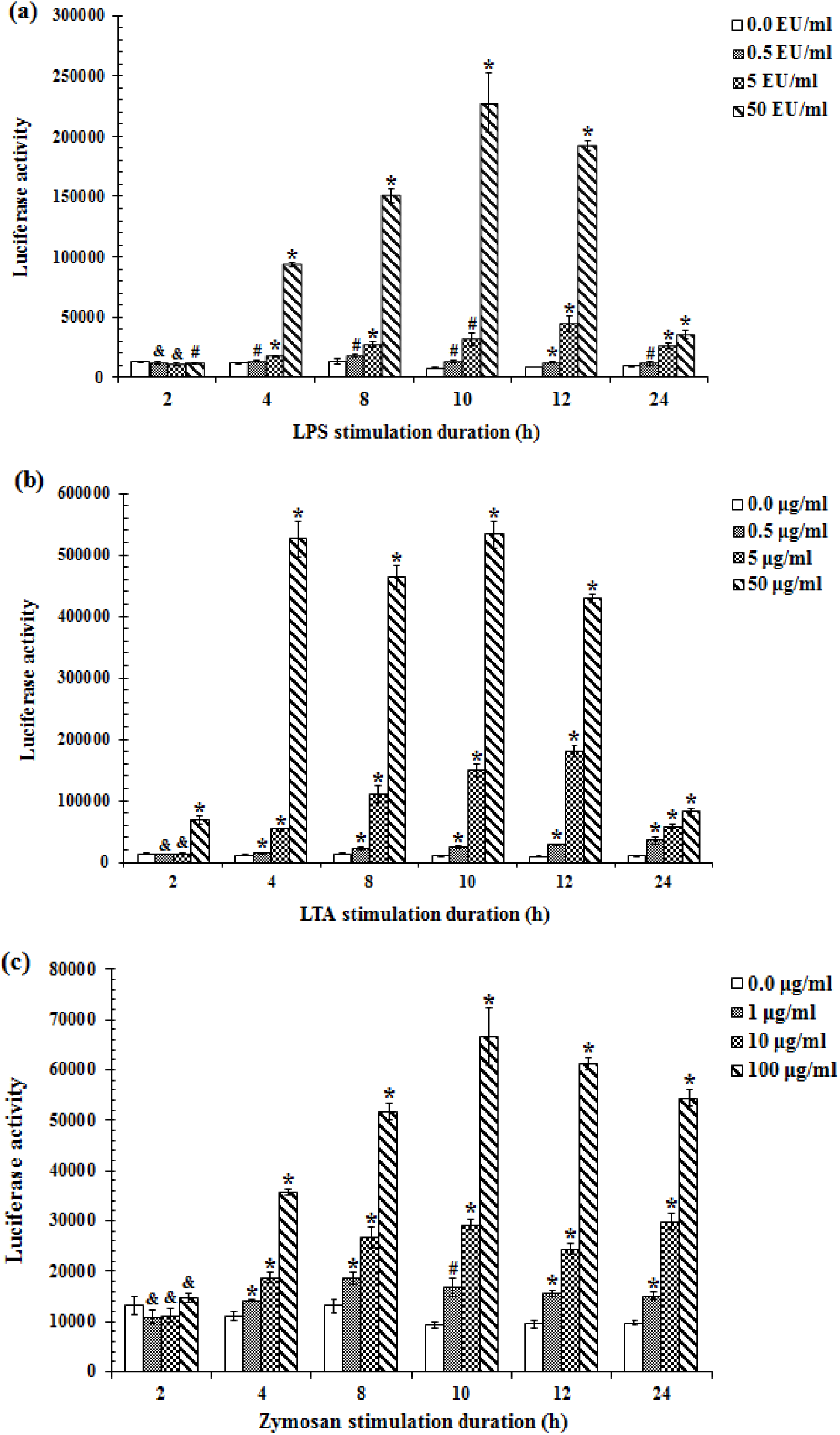
The NF-κB activity levels of RAW 264.7 cells at a density of 10×10^5^ cells/ml after stimulation with LPS (a), LTA (b), and zymosan (c) for different times (*n*=4). & *P*>0.05 vs. the negative control (e.g., 0 EU/ml, 0 μg/ml) # *P*<0.05 vs. the negative control * *P*<0.01 vs. the negative control

### 3.4. The dose-effect relationships of LPS in activating NF-κB in RAW 264.7 cells at different cell densities and passages

The results of this experiment are presented in Figure 4. The data of Figure 4a show that at different cell densities, LPS had a dose-effect relationship with NF-κB activity to a certain extent. The dose-response curve was relatively flatter for low cell density (5×10^5^ cells/ml) than for the medium and high cell densities (10×10^5^ and 15×10^5^ cells/ml, respectively). However, LPS had a relatively narrower concentration range (32-128 EU/ml) for maintaining maximum NF-κB activity at the high cell density (15×10^5^ cells/ml) compared with that (32-512 EU/ml) at the medium cell density (10×10^5^ cells/ml). Besides, the NF-κB activity at the high cell density (15×10^5^ cells/ml) was much more susceptible to the “edge effect” of plates in the culture, which might lead to a decrease in NF-κB activity at that density stimulated by the 512 EU/ml of LPS. The data of Figure 4b show that at different cell passages with a density of 10×10^5^ cells/ml, LPS also had a stable dose-effect relationship with NF-κB activity. The dose-response curves at different cell passages were relatively parallel. These results indicate that LPS had the best and most stable dose-effect relationship with NF-κB activity at a cell density of 10×10^5^ cells/ml.

**Fig. 4.**
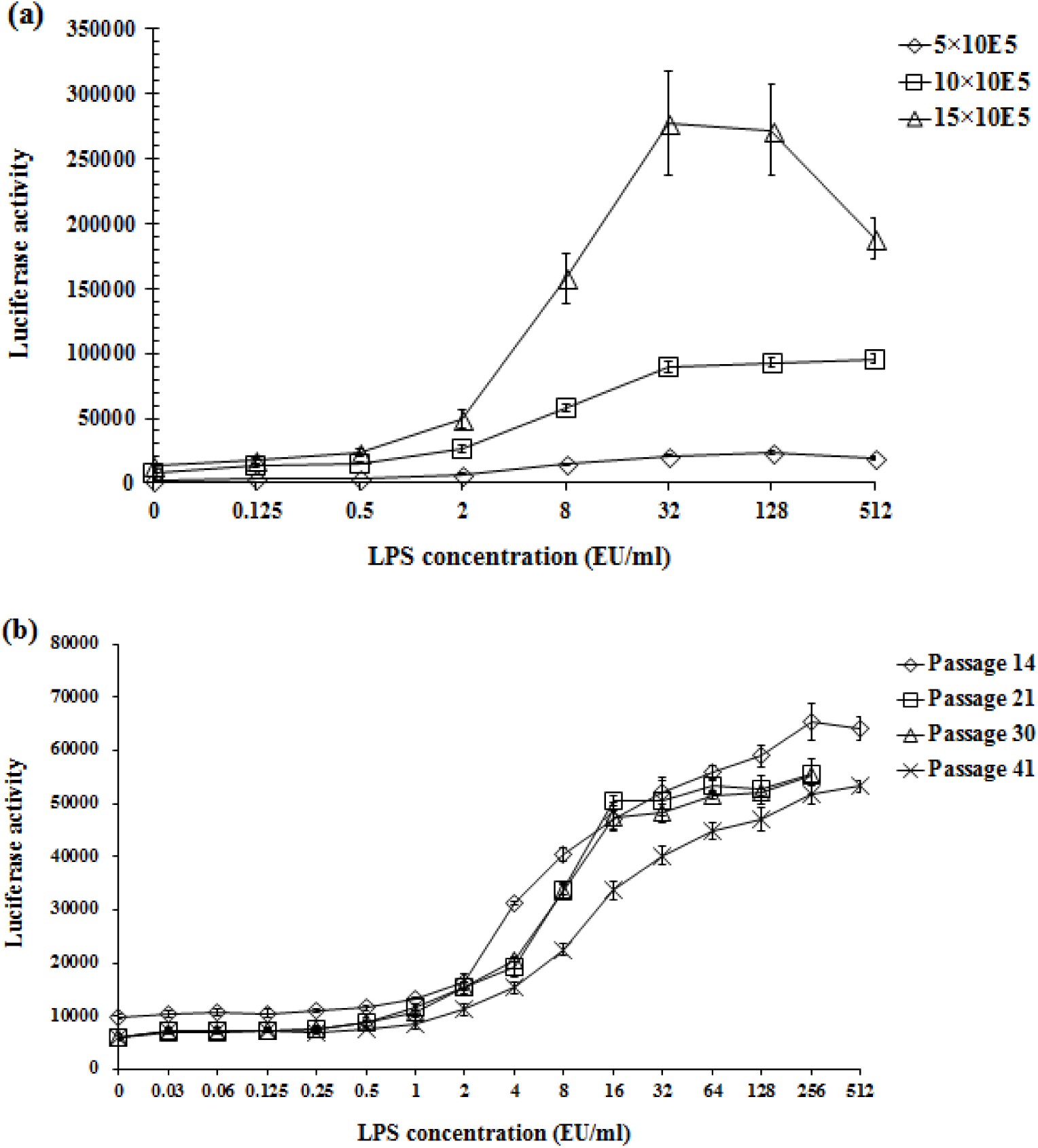
The NF-κB activity levels of RAW 264.7 cells at 5×10^5^, 10×10^5^, and 15×10^5^ cells/ml (a); and at passage 14, 21, 30, and 40 with a density of 10×10^5^ cells/ml (b) after stimulation with LPS for 10 h (*n*=4).

### 3.5. The limits of detection (LODs) and dose-response curves of pyrogen-induced NF-κB responses

The results of this experiment are presented in Figure 5. The data show that the LODs of the NF-κB responses to LPS, LTA, and zymosan were 0.03 EU/ml (*P*=0.005), 0.001 μg/ml (*P*=0.021), and 1 μg/ml (*P*=0.033), respectively. The activity of NF-κB increased in a dose-dependent manner in response to the pyrogen concentration. These results indicate the good sensitivity and dose-effect relationship of this method for detecting pyrogens.

**Fig. 5.**
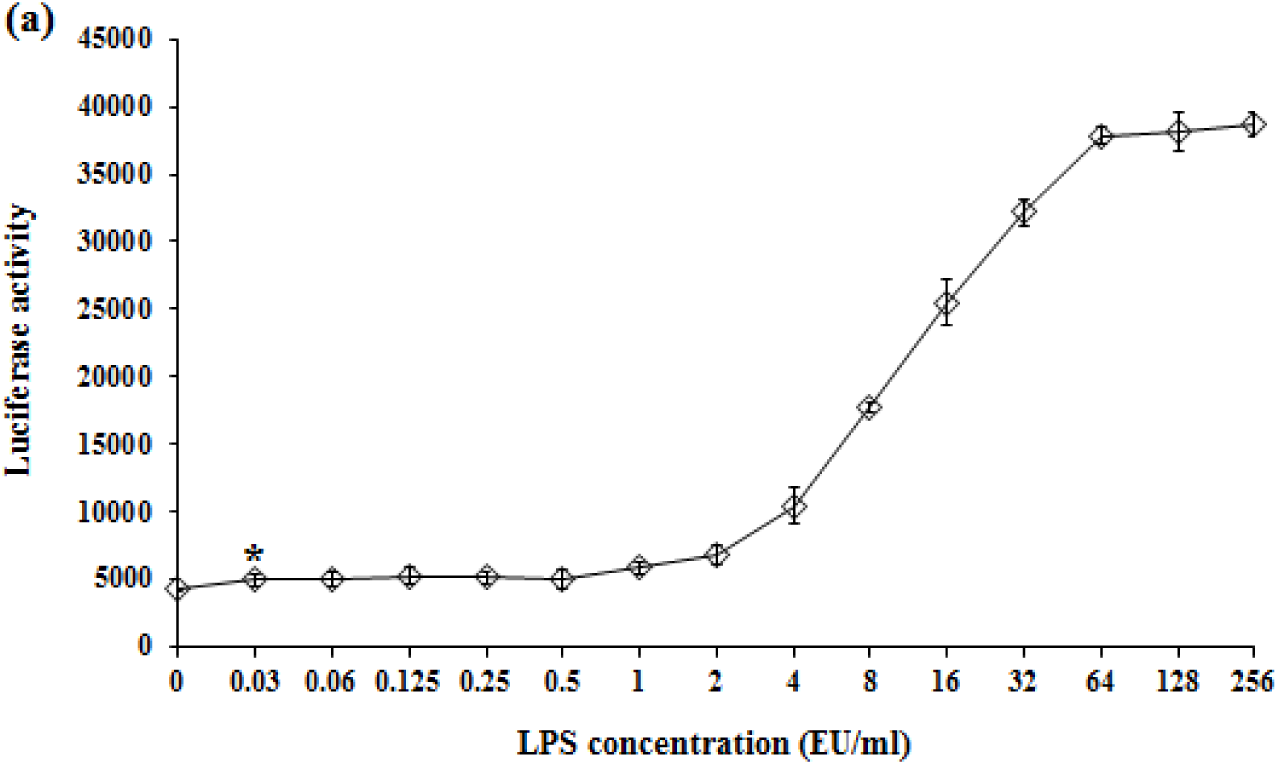
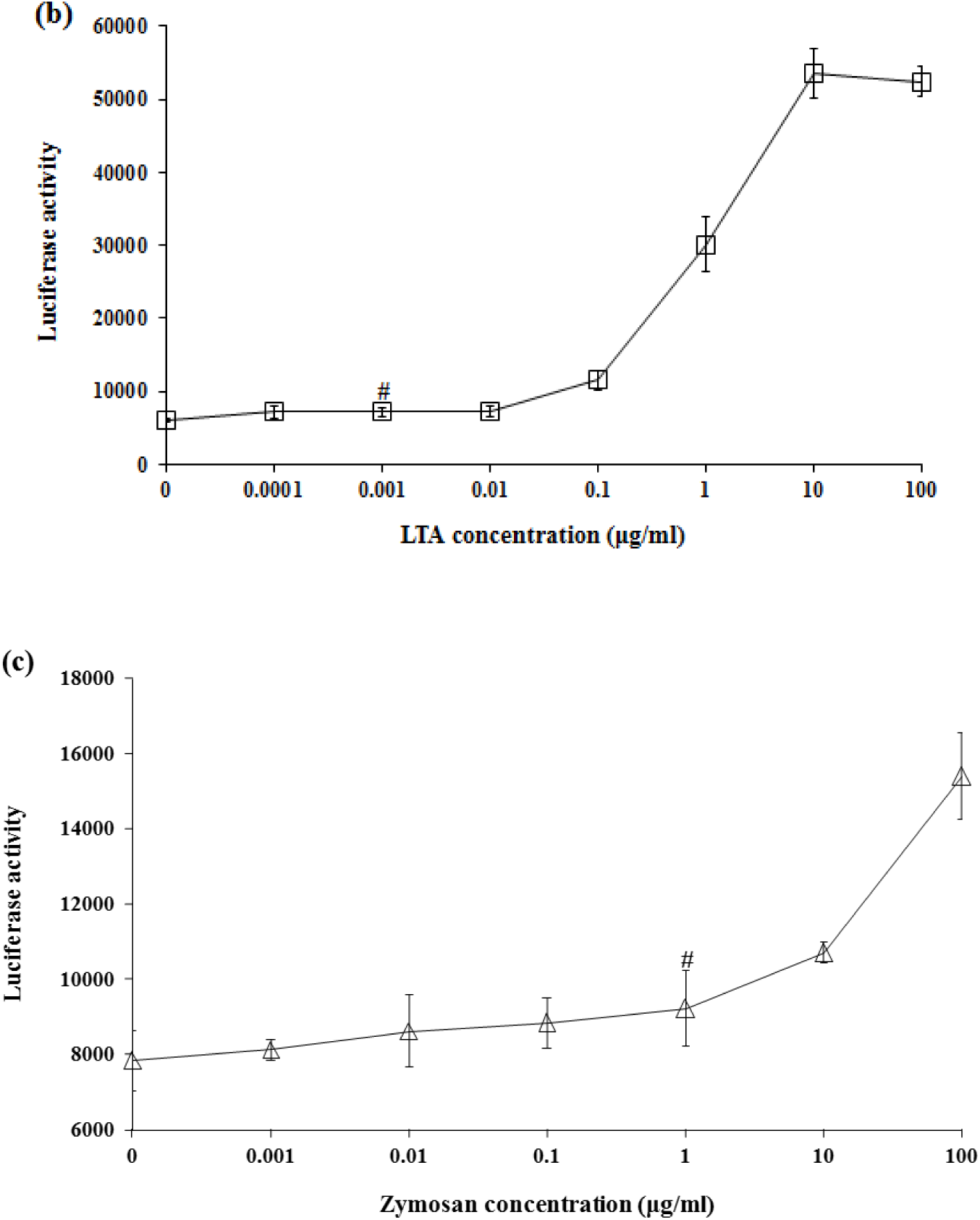
The NF-κB activity levels of RAW 264.7 cells at a density of 10×10^5^ cells/ml after stimulation with a series of concentrations of LPS (a), LTA (b), and zymosan (c) for 10 h (*n*=4). # *P*<0.05 vs. the negative control (e.g., 0 EU/ml, 0 μg/ml) * *P*<0.01 vs. the negative control

### 3.6. Precision and accuracy of the RGA for detecting LPS in the laboratory

Four samples of LPS, whose expected concentrations were 1.5 (sample 1), 3.0 (sample 2), 6.0 (sample 3), and 12.0 (sample 4) EU/ml, respectively, were detected by the RGA. The results are presented in Table 1. The data show that the measured concentrations of LPS in the samples were 1.761±0.126 (sample 1), 3.241±0.101 (sample 2), 5.906±0.390 (sample 3), and 11.028±1.760 (sample 4) EU/ml, respectively, and the intraassay and interassay coefficients of variation (CVs) were generally less than 13% and 16%.

The results are presented in Figure 6. The data show a good linear relationship between the expected and measured concentrations in the tested LPS concentration range (1.5-12.0 EU/ml), suggesting that the method also had good accuracy.

**Table 1.**
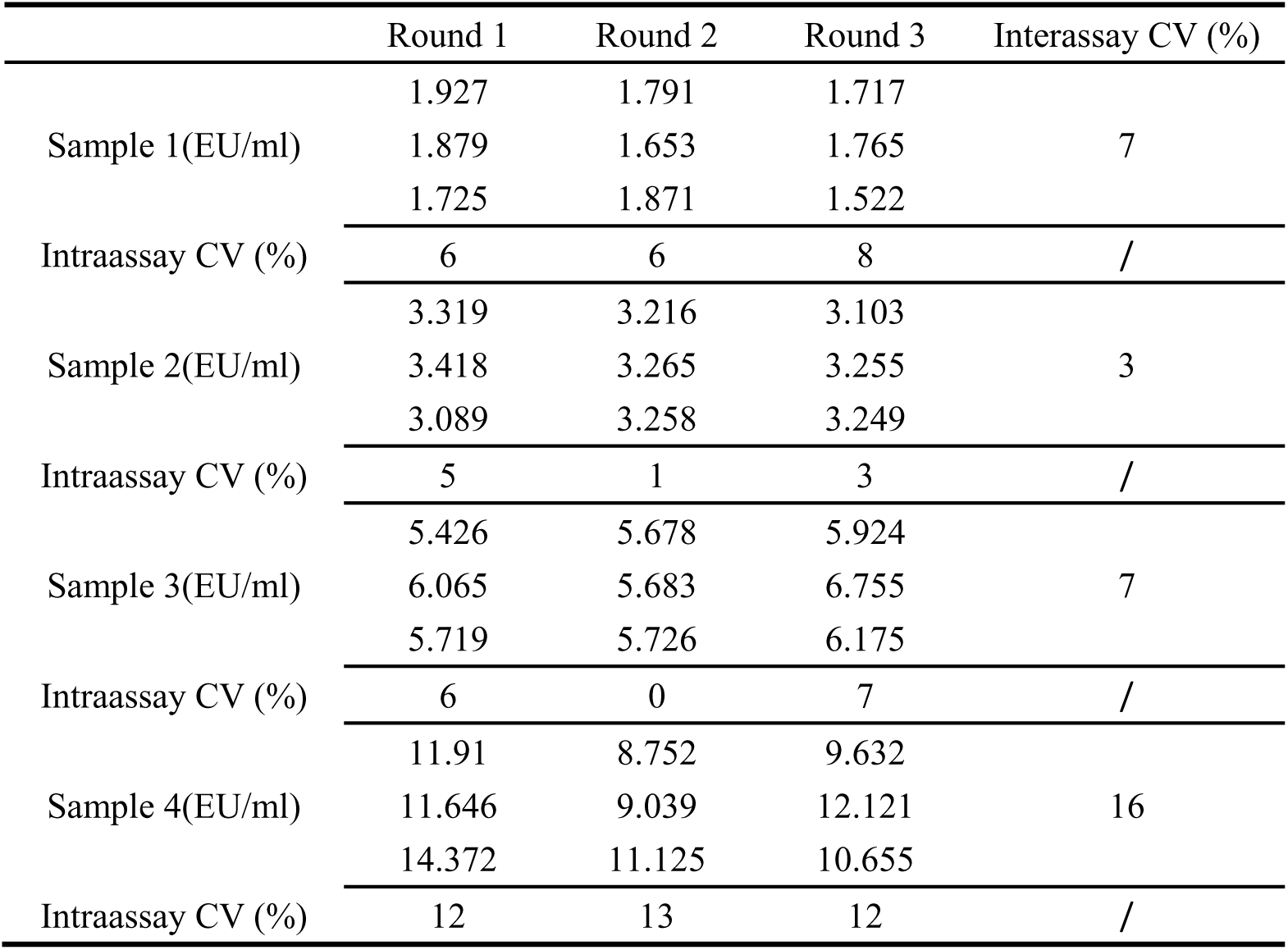
Precision of the RGA for detecting LPS

**Fig. 6.**
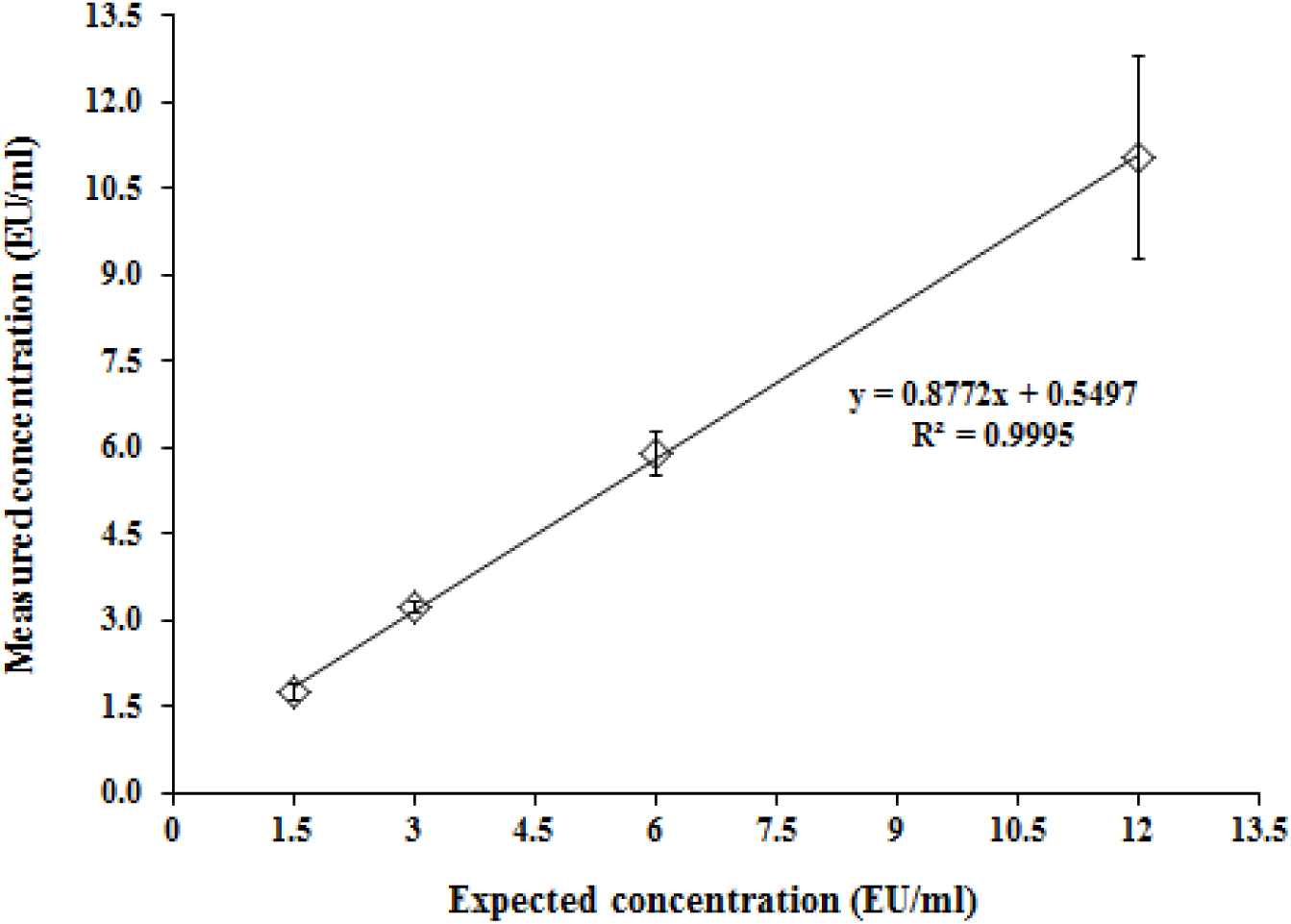
Linear regression analysis between the expected and measured LPS concentrations.

### 3.7. Application of the RGA to drugs

Drugs within the maximum valid dilution (MVD) pass the interference assay when the spike recovery is within the range of 50-200%. The recovery concentration of LPS in the drugs was 4.0 EU/ml, and the LOD of LPS used in the experiment was 0.125 EU/ml. The results are presented in Table 2, and they indicate that the method has potential for various applications.

**Table 2.** Recovery of LPS spike in drugs measured by the RGA

| Drug | Fold-dilution | NF-κB response |  |
| --- | --- | --- | --- |
|  |  | Spike recovery (%) | Interference |
| Nivolumab injection | 16 | 121 | no |
| Rituximab injection | 8 | 125 | no |
| Bevacizumab injection | 16 | 161 | no |
| Etanercept solution for injection | 168 | 105 | no |
| *Haemophilus influenzae* type b conjugate vaccine | 400 | 74 | no |
| 23-Valent pneumococcal polysaccharide vaccine | 400 | 75 | no |
| Group A and group C meningococcal conjugate vaccine | 8000 | 85 | no |
| Basiliximab for injection | 64 | 70 | no |

Spike recovery=100%(the mean concentration of endotoxin detected in the diluted solution containing the added endotoxin - that detected in the diluted solution)/the added endotoxin.

### 3.8. Comparison between the RGA and BET

Drugs were tested at their MVD, each of which was calculated as the endotoxin limit concentration in EU/ml divided by the LOD (in this case, 0.5 EU/ml). Each drug presented five blinded spikes, two of which were defined as negative, meaning they were below 0.5 EU/ml (0 and 0.25 EU/ml), while three were positive (0.5, 1.0, and 2.0 EU/ml). These spikes were tested by both the RGA and BET. After the interference test was passed, the samples were classified by a so-called prediction model (PM), which classified the samples as negative (N) when the mean concentration of endotoxin equivalents in each of the sample replicates calculated by the endotoxin standard curve was less than the contaminant limit concentration specified for the samples. The samples were otherwise classified as positive (P). Within-laboratory reproducibility was calculated as the proportion of samples classified identically in three independent runs. Reproducibility between methods was calculated as the proportion of samples classified identically. Sensitivity was defined as the probability of correctly classifying positive samples, and specificity was defined as the probability of correctly classifying negative samples. The results are presented in Table 3.

**Table 3.** Classification of samples by the RGA and BET

| Drug | Spike<br>(EU/ml) | Truth | RGA |  |  | BET |  |  |
| --- | --- | --- | --- | --- | --- | --- | --- | --- |
|  |  |  | Round | Round | Round | Round | Round | Round |
|  |  |  | 1 | 2 | 3 | 1 | 2 | 3 |
| *Haemophilus influenzae* type b conjugate vaccine | 0 | N | N | N | N | N | N | N |
|  | 0.25 | N | N | N | N | N | N | N |
|  | 0.5 | P | **N** | **P** | **P** | **N** | **N** | **N** |
|  | 1 | P | P | P | P | P | P | P |
|  | 2 | P | P | P | P | P | P | P |
| 23-Valent pneumococcal polysaccharide vaccine | 0 | N | N | N | N | N | N | N |
|  | 0.25 | N | N | N | N | N | N | N |
|  | 0.5 | P | P | P | P | **N** | **N** | **N** |
|  | 1 | P | P | P | P | P | P | P |
|  | 2 | P | P | P | P | P | P | P |
| Sensitivity (%) |  |  | 94 (17/18) |  |  | 67 (12/18) |  |  |
| Specificity (%) |  |  | 100 (12/12) |  |  | 100 (12/12) |  |  |
| Within-laboratory reproducibility (%) |  |  | 90 (27/30) |  |  | 100 (30/30) |  |  |
| Reproducibility between assays (%) |  |  | 87 (52/60) |  |  | / |  |  |

False classifications are in bold type.

The BET assay, in principle, is a physicochemical binding assay. Protein components in products often interfere with the binding of LPS to the limulus agents, leading to the fact that the interference of the product is mostly manifested as inhibition. Even if the interference of the product at a small dilution is eliminated, false-negative results are more likely to occur in the BET, as shown by the sensitivities of the BET (67%) and RGA (94%). However, the accuracy of the BET will be affected when the interference of the product at a large dilution is eliminated. The RGA can overcome the above problems of the BET, and the results of the RGA should be more accurate.

## 4. Discussion

Currently, three pyrogen tests, including the RPT, BET, and MAT, have been adopted by pharmacopeias [3,4]. The RPT involves the use of animals *in vivo*, does not correspond with the 3Rs principle, has variations in response depending on many factors, and is expensive. The BET is a physicochemical test for detecting the LPS of gram-negative bacteria, not a functional activity test [33]. The MAT is mainly based on the use of monocytes and macrophages involved in the febrile response and can overcome those problems; however, it often needs large amounts of human blood and its convenience needs to be improved. The representativeness of using a single proinflammatory cytokine as the pyrogenic marker is also limited in theory.

Compared with other sources of monocytes and macrophages, such as whole blood [34-36] and peripheral blood monocytes [37-39], monocytic and macrophage cell lines are more stable and easier to use [40-43]. RAW 264.7 cells are mouse-derived macrophages that contain a variety of receptors involved in the immune response, as described above, and can react with various pyrogens. The results of this study also confirm that pyrogens from different sources can activate an NF-κB reporter gene when it is transfected into RAW 264.7 cells.

Most existing MATs often use various proinflammatory cytokines (e.g., IL-6, IL-1β, and TNF-α) as pyrogenic markers [6,44]. Although this approach has a certain rationality, pyrogens stimulate the body via different mechanisms to induce immune cells to secrete different proinflammatory factors. Our previous study found differences between the secretion of IL-6 and IL-1β even from the same cryopreserved or fresh pooled human whole blood [45,46]. However, the synthesis and secretion of various proinflammatory factors mostly involve the activation of NF-κB; thus, it is more reasonable to use NF-κB as a representative pyrogenic marker. The results of this study confirm that the activation of NF-κB by stimulation with different pyrogens, including LPS, LTA, and zymosan, showed a good correlation with the secretion of proinflammatory factors (e.g., IL-1β, IL-6, and TNF-α) induced by those pyrogens. Additionally, the time-effect relationships of the induced NF-κB activation were similar among the pyrogens, which might also verify that NF-κB is a central signaling molecule that mediates the fever reaction induced by pyrogens. It has been reported that RAW 264.7 and THP-1 cells have similar detection limits for LPS and LTA [40]. We also found that fresh pooled human whole blood and RAW 264.7 cells had similar reactivities to LTA and zymosan [46].

This study verifies the feasibility of the novel RGA for pyrogen detection in the laboratory. We plan to organize validation of the method in different laboratories. In conclusion, this study establishes a novel bioassay for pyrogen detection using RAW 264.7 cells transfected with a NF-κB reporter gene as a pyrogenic marker. This method can be used to detect multiple pyrogens; is sensitive, stable, and accurate; and can be applied widely.

## Acknowledgement

We thank Zean Yang and Yusheng Pei for their invaluable technical assistance.

